# Oatmeal and wheat flour as the sources of thyroid peroxidase (TPO), lipoxygenase (LOX) and xanthine oxidase (XO) modulators potentially applicable in the prevention of inflammatory thyroid diseases

**DOI:** 10.1101/2023.06.05.543703

**Authors:** Ewa Habza - Kowalska, Katarzyna Piwowarczyk, Jarosław Czyż, Urszula Gawlik - Dziki

## Abstract

Despite the widespread potential pro-health effects of ferulic acid (FA), their interference in the progression of thyroid dysfunction has mainly remained unresolved. Here, we combined in vitro enzyme studies with the in vitro cellular approach to investigate the potential of main dietary sources of FA - the oatmeal (OM) and wheat flour (WF) compounds for the prophylactics of inflammatory thyroid diseases. Potentially bioaccessible OM and WF compounds activated thyroid peroxidase (TPO), while inhibiting the activity of lipoxygenase (LOX) and xanthine oxidase (XO). Isobolographic studies revealed cooperation between them. Relatively strong inhibitory activity of bioaccessible OM compounds on LOX activity correlated with their cytostatic and pro-invasive effects in thyroid cancer model in vitro. These data indicate the potential of OM and WF products for the prophylactics of inflammatory thyroid diseases (incl. hypothyroidism). However, it should be considered with care, especially in the context of the oncological status of the patient.

## 1. Introduction

Inflammation is a natural process that occurs in tissues as a defensive mechanism against tissue injury or microbial infection. However, chronic inflammation is often associated with the pathogenesis and progression of autoimmune, cardiovascular and neurological diseases (Jaismy et al., 2018). It also contributes to the autoimmune destruction of the thyroid gland (Danailova et al., 2022). Development of thyroid diseases, like of many other autoimmune disorders, is believed to result from the combinations of versatile stimuli, including environmental, lifestyle, and genetic factors. For instance, regular consumption of “proinflammatory” food can result in the intestinal inflammation that spreads to other organs in the body, eventually contributing to the development of thyroid diseases, incl. hypothyroidism and cancer (Kochman et al., 2021).

Thyroid malfunctions prevalently occur in women. Overt hyperthyroidism results from an overproduction of thyroid hormones and is primarily related to Graves’ disease. In turn, hypothyroidism results from a deficiency of thyroid hormones, and its most prevalent syndrome is Hashimoto’s thyroiditis (Xu et al., 2019). Several clues indicate the involvement of redox enzymes in its etiopathology. For instance, a deficiency or abnormal function of TPO can lead to impaired thyroid hormone synthesis, resulting in congenital hypothyroidism. TPO plays a key role in catalyzing the oxidation of iodide, which is necessary for the iodination of tyrosyl residues in thyroglobulin (TG) - a process known as organification. Additionally, TPO is responsible for the oxidative coupling of iodothyronine residues to form the hormones T4 and T3. In hypothyroidism, TPO acts as an autoantigen, prompting the generation of circulating autoantibodies and thyroid inflammation in patients with Hashimoto’s thyroiditis (Williams, 2008).

Inflammatory etiopathology of thyroid diseases implies the involvement of inflammatory mediators/enzymes in their development. For instance, cyclooxygenase (COX) and lipoxygenase (LOX) are responsible for a wide range of physiological and pathophysiological responses (Yao et al., 2015). Lipoxygenases (LOXs) are responsible for the oxygenation of polyunsaturated fatty acids, such as arachidonic acid, to bioactive lipids, including leukotrienes and hydroxyeicosatetraenoic acids (HETEs). LOX products are involved in a wide range of physiological processes, including inflammation, immune responses, and the proliferation of cancer cells, thus linking the inflammation with carcinogenesis (Yao et al., 2015; Zabiulla et al., 2022). While the participation of COXs and LOXs in inflammatory processes and in oxidative stress induction is well documented, the participation of xanthine oxidase (XO) in these processes has recently gained an increasing interest. Its activity can result in the conversion of superoxide radicals into hydroxyl radicals, which exacerbate inflammatory responses and contribute to the development of the cytokine storm syndrome (CSS) (Pratomo et al., 2021). Whereas several studies reported on the inhibitors that can concomitantly affect the COX and LOX activity (Jaismy et al., 2018), only few studies have been focused on the substances that concomitantly affect XO/LOX activity. Given the possible interrelations between the inflammation and oxidative stress in hypothyroidism, there is also a need to identify and isolate LOX/XO inhibitors that would concomitantly activate TPO (Zabiulla et al., 2022).

We have previously shown that pure polyphenolic substances, such as ferulic acid (FA), can enhance TPO activity *in silico* (Habza-Kowalska, Kaczor, Żuk, Matosiuk, & Gawlik-Dziki, 2019; Habza-Kowalska, Kaczor, Bartuzi, Piłat, & Gawlik-Dziki, 2021). On the other hand, the interference of FA-rich plant products with the activity of LOX and XO has not been addressed so far. In the current study, we (i) identified wheat flour (WF) and oat flakes (oatmeal; OM) as the products relatively rich in FA. Then, we estimated the interference of bioaccessible OM and WF compounds with TPO, LOX and XO activities to assess their potential in diet supplementation for the patients with hypothyroidism. Finally, (iii) using an in vitro cellular approach, we addressed the potential consequences of their activity for the patients with thyroid carcinoma. To the best of our knowledge, this work is the first to consider the activity of diet TPO/LOX/XO modulators in the context of their side effects.

## 2. Materials and Methods

### 2.1. Chemicals

Sucrose (α -D-glucopyranosyl-(1→4)- β -D-fructofuranoside), Tris (1,3-Propanediol-2-amino-2-hydroxymethyl), KCl, NaCl, MgCl_2_, 90% ethanol, NaOH, guaiacol (2-Methoxyphenol), H_2_O_2_ (Hydrogen Peroxide), ABTS (2,2′-azinobis-(3-ethylbenzothiazoline-6-sulfonic acid), lipoxygenase (LOX), linoleic and ferulic acids, xanthine oxidase (XO), xanthine, pancreatin, pepsin, bile extract, phosphate buffered saline - pH 7.2 (PBS) were purchased from Sigma-Aldrich Company (Poznan, Poland). All other chemicals were of analytical grade.

### 2.2. Material

Porcine thyroid glands were purchased at a local slaughterhouse (Lublin, Poland) and stored in -20°C until used. Oatmeal (Plony Natury, Poland) was purchased in local supermarket in Lublin, Poland. Flour from common wheat (cv. Batuta) was purchased in the local mill (Lublin, Poland).

### 2.3. Preparation of EtOH, raw and digested extracts of polyphenolic plant sources

For ethanol extraction, 1.5 g of individual raw materials were homogenized with 15 mL of 50 % ethanol, samples were shaken for 30 min in the room temperature and then centrifuged for 15 min at 4000 RPM. The extraction procedure was repeated twice. The final samples were brought to 50 ml with 50% ethanol to reach the extract concentration of 30 mg/mL. Extracts were diluted to concentration 0.3 mg/mL. 50% ethanol extracts were used as a control for enzymatic tests and determination of antiradical potential. For *in vitro* tests, PBS extracts (containing potentially bioavailable free compounds) were used to eliminate the potential effect of ethanol. It was prepared analogously to the ethanol extract. *In vitro* digestion of raw materials was performed according to the modified procedure proposed by Minekus et al. (Minekus et al., 2014). 0.5 g of row material was mixed with 5 ml of simulated saliva fluid (SSF) containing 0.22 mg of α-amylase and 200 μl of 5M HCl) and incubated for 2 min in 37°C. Then, 250 μL of SGF (750 μL of pepsin mixed with 50 mL SGF) was added and incubated for 2 hours in temperature 37°C to simulate gastric digestion, followed by the application of 1.2 ml of 0.1M NaHCO_3_ (pH 6.0) and 750 μL of 1M NaHCO_3_. Finally, 600 μL of SIF (0.05 g of pancreatin, 0.3 g of bile extract in 35 ml of SIF; pH 6,0) as applied, the sample was incubated for 1 hour in 37°C and centrifuged at 4500 rpm for 15 minutes. The supernatant was stored at -20°C until further analyses. 0.5 ml of distilled water was used instead of plant source to obtain the control sample.

### 2.4. Determination of total phenolics content (TPC)

TPC analyses were carried out with the protocol of Singleton and Rossi (Singleton & Rossi, 1965) adopted for microplate reader (Epoch 2 Microplate Spectrophotometer, BioTek Instruments, Winooski, Vermont, USA). Ten microliters of extract, 10 μL of water and 40 μL of Folin-Ciocalteau reagent were diluted in water at the ratio of 1:5. After 3 min, 250 μL of 10% sodium carbonate was added and the solution was thoroughly mixed. 50% ethanol or digested control (H_2_O) sample was used as the standard. The absorbance was measured at 725 nm after 30 minutes of incubation and normalized against the standard. The concentration of phenolic compounds was read from the standard curve determined for gallic acid and expressed as gallic acid equivalent (GAE) in mg/g DW.

### 2.5. *In vitro* antioxidant capacity assay

ABTS radical scavenging activity was prepared according to Re et al. (1999) (Re et al., 1999) with some modifications. 250 μl ABTS was mixed with 10 μl of the sample (ethanol and GD extracts) and measured at the wavelength 724 nm using UV/Vis microplate reader (Epoch 2 Microplate Spectrophotometer, BioTek Instruments, Winooski, Vermont, USA). after 15 min of incubation in RT. ABTS discoloration was calculated as follows:

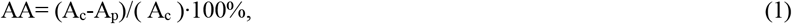

where:

A_c_ – the absorbance of control, A_p_ – the absorbance of extract

DPPH radical scavenging activity was measured according to (Brand-Williams et al., 1995) with some modifications. 250 μl of DPPH solution was mixed with 10 μl of the extract (3 mg/ml) and measured at 517 nm using UV/Vis microplate spectrophotometer (BioTek, Model Epoch2TC, Winooski, Vermont, USA) after 15 minutes of incubation in RT. The inhibition percentage of DPPH discoloration was calculated as in (1).

### 2.6. Thyroid peroxidase (TPO) activity assay

The assays was prepared according to (Jomaa, 2015), with some modifications. Detailed description is provided in the publication by Habza-Kowalska et al. (Habza-Kowalska, Gawlik-Dziki, et al., 2019). TPO activity was calculated using the formula below:

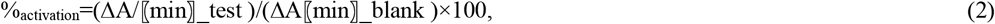

Where: ΔA/min test is the linear absorbance change per minute of the test material and ΔA min blank is the linear change in absorbance per minute of blank. The mode of enzyme activation/inhibition estimated with the Lineweaver-Burk plot.

### 2.8. Inhibition of lipoxygenase (LOX) activity

LOX activity was analyzed using the protocol proposed by Axelrod et al., (1981) adopted for microplate reader (Epoch 2 Microplate Spectrophotometer, BioTek Instruments, Winooski, Vermont, USA). Detailed description can be found in the publication by Habza-Kowalska et al. 2019 (Habza-Kowalska, Gawlik-Dziki, et al., 2019). LOX inhibitory activity was calculated using the formula (3). The mode of inhibition of the enzyme was performed using the Lineweaver-Burk graph. The EC_50_ values were calculated from the fitted models of dose-dependence and given as the concentration of the tested compound that gave 50% of the maximum inhibition.

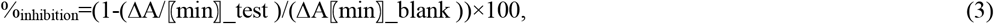

Where: ΔA/min test is the linear absorbance change per minute of the test material and ΔA min blank is the linear change in absorbance per minute of blank.

### 2.9. Inhibition of xantine oxidase (XO) activity

XO activity was measured according to Sweeney et al. (Sweeney et al., 2001) with some modifications: 30 μL of the sample was diluted in 110 μL of 1/15 M/L phosphate buffer (pH 7.5), and 20 μL of enzyme solution (0.01 U/ml in M/15 phosphate buffer). After the preincubation at 30°C for 10 min, the reaction was started by adding 140 μL of 0.15 mM/L xanthine solution. The absorbance (295 nm) was measured every minute for 3 min. XO inhibitory activity was calculated using the formula (3). Inhibitory activity was expressed as EC_50_ (efficient concentration) i.e. the amount of sample needed to inhibit XO to 50% of initial activity.

### 2.10. Isobolographic analysis

Isobolographic analyses were performed according to (Chou, 2006). A detailed description can be found in the publication by Habza-Kowalska et al. (Habza-Kowalska, Gawlik-Dziki, et al., 2019).

### 2.11. In vitro approach

#### 2.11.1. Cell culture

Human thyroid cancer B-CPAP (ACC 273, DSMZ-German Collection of Microorganisms and Cell Cultures GmbH) and 8505C cells (Sigma No. 94090184) were cultured in the standard conditions in RPMI 1640 and DMEM/F12 HAM, respectively (Sigma No. <u>D8437</u>) supplemented with 10% heat-inactivated fetal bovine serum (FBS; Gibco, No. A3840402) and 1% Antibiotic-Antimycotic Solution (Merck, No. A5955). For each experiment, the cells were harvested with Ca^2+/^Mg^2+^-free 0,25% trypsin/EDTA/PBS solution (Gibco No. <u>25200072</u>), counted in Z2 particle counter (Beckman Coulter) and seeded into multi-well tissue culture plates (Falcon®). Cells were exposed to extracts administered at the concentration of 0.01, 0.05, 0.1, 1 and 3% in culture medium (corresponding to 1.5, 7.5, 15, 150 and 450 g of the product/75 kg body mass) for 48 hours.

#### 2.11.2. Proliferation and viability tests

For the analyses of cell viability and proliferation, B-CPA and 8505C cells were seeded into 12–well cell culture plates (Corning®Costar®) at the density of 3 and 2.5×10^4^ cells/well and cultivated for 24 hours before the administration of the extract in a fresh culture medium. Cell viability was estimated with EtBr/FDA assay 48 hours after extract administration. Proliferation was estimated with Coulter counter 48 hours after the administration of extracts (Ryszawy et al., 2019).

### 2.11.3. Wound healing assay

B-CPAP and 8505C cells were seeded in 12-well plates at the density of 400 cells/mm^2^. After 24 hours, the extracts were added along with the medium at the concentration of 0.1 and 1%. After the next 48 hours, a wound was made in the center of each well with a clean tip and 16 wound pictures were registered immediately afterwards and 24 hours thereafter to calculate the %age of wound coverage. The analyses were performed at 37°C and 5% CO2 using a Leica DMI6000B fluorescence microscope.

### 2.11.4. Immunofluorescence

For immunofluorescence studies, B-CPAP and 8505C cells were seeded into 12-well plates on UVC-sterilized coverslips at the density of 3 and 2.5×10^4^ cells/well, respectively, cultured for 24 hours and processed in the presence/absence of the extracts administered at the concentrations given in the text. Then, they were fixed with 3.7% formaldehyde followed by 0.1% Triton X-100 permeabilisation (Pudełek et al., 2020) and non-specific binding sites were blocked with 3% BSA (Invitrogen, No. 37525; 30 min. in 37°C). After washing with 2% PBS, the mouse monoclonal anti-vinculin IgG (with 1% BSA, Sigma no. V9131) was applied for 45 min. Then, the specimens were washed before the application of the mixture of AlexaFluor488-conjugated goat anti-mouse IgG (ThermoScientific No. A-11029), AlexaFluor546-conjugated phalloidin (Invitrogen, No. A22283; for F-actin visualization) and Hoechst 33258 (Sigma; for DNA staining) for 45 min. Finally, specimens were mounted in Agilent Dako mounting medium (Agilent Dako; No. S3023). Images were acquired with Leica DMI6000B fluorescence microscope equipped with DFC360FX CCD camera. For better clarity, the raw images were additionally processed (linear contrast adjustment and background subtraction) in ImageJ software.

### 2.11.5. Statistical analysis

All data were expressed as mean +/-SEM from at least three independent experiments (n = 3). The statistical significance was tested with one-way ANOVA followed by post-hoc Dunnett’s or Tukey’s comparison for variables with a non-normal (tested with Levene’s comparison) and normal distribution, respectively. Statistical significance was shown at p < 0.05.

## 3. Results and Discussion

### 3.1. Determining Ferulic Acid Content in Natural Plant-Based Food Sources Using Phenol Explorer Database

Previously, we have shown versatile effects of purified polyphenols and their natural sources (plant extracts) on the activity of TPO and LOX, as the enzymes involved in thyroid diseases (Habza-Kowalska, Kaczor, et al., 2019; Habza-Kowalska, Gawlik-Dziki, et al., 2019). For instance, ferulic acid (FA) has been shown to activate TPO (Habza-Kowalska et al., 2021), which indicates its potential in the prophylactics of hypothyroidism. Ferulic acid is a naturally occurring phenolic compound of plant-based foods, which displays antioxidant, anti-inflammatory, and anticancer properties (Nile et al., 2016). The content of FA varies between food sources, which points to the necessity for the identification of FA-rich products. Based on our preliminary studies on FA activity, we first concentrated on the identification of natural sources of FA (Patel et al., 2013). Phenol Explorer (http://phenol-explorer.eu/) database has been developed to provide comprehensive information on the content of phenolic compounds in foods. Previously, it allowed to identify the sources of FA (grains, fruits, and vegetables), which can easily be incorporated into a healthy diet to increase the intake of this bioactive compound (Ramli et al., 2017). Using this database, we pinpointed oatmeal (OM) and wheat flour (WF) as the rich FA sources.

Detailed studies on the qualitative/quantitative profile of phenolic acids in OM and WF used in this study have recently been published (Soycan et al., 2019; Buczek et al., 2023). These data indicate that WF and OM are rich sources of this compound. As shown in the she summary of these data (Table 1), WF contains almost 3-fold higher FA content than OM. Therefore, in the following experiments we focused on the activity of the OM and WF extracts. These extracts were subjected to simulated gastrointestinal digestion (GD) and their activity was compared to the commonly studied 50% EtOH WF and OM extracts.

**Table 1.**
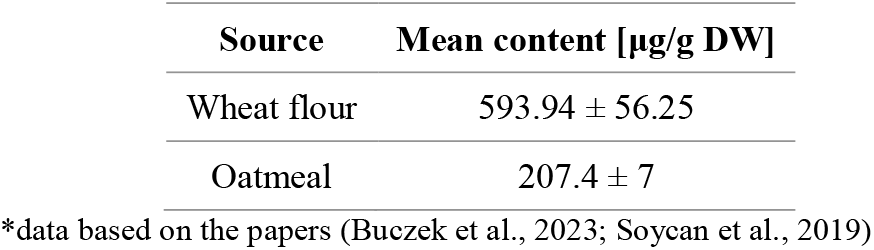
Ferulic acid content*

## 3.2. TPC and anti-oxidative activity of WF and OM extracts

Total phenolic content (TPC) is a widely used predictor of the potential antioxidant and anti-inflammatory activity of plant extracts. Our preliminary experiments were performed to estimate the basic parameters of EtOH and GD extracts from wheat flour (WF) and oatmeal (OM). TPC estimated for EtOH WF extracts reached 14.25 ± 0.71 mg GAE/g dry weight (Table 2). It was considerably higher than that estimated for EtOH OM extracts (5.66 ± 0.28 mg GAE/g dry weight), but lower than the values reported by Călinoiu & Vodnar (2020) for MetOH WF and OM extracts (WF: 39.61 ± 0.51 mg GAE/100 g dry weight; OM: 25.15 ± 0.45 mg GAE/100 g dry weight). These differences could be attributed to different sample preparation, and the type and quality of the wheat flour and oatmeal used. For instance, TPC of water extracts from WF can range from 211.55 to 1393.27 μg GAE/g (Tian et al., 2021;Yu & Beta, 2015). The EtOH extraction method can also be less effective than the others, even if it is commonly used in the preparation of para-pharmaceutics. Accordingly, gastrointestinally digested (GD) WF and OM extracts were characterized by higher TPCs (18.60 ± 0.93 and 11.59 ± 0.58 mg GAE/g dry weight, respectively) than their EtOH counterparts. This observation indicates that GD effectively releases phenolic compounds from WF and OM and justifies further focus on the properties of GD extracts.

**Table 2.**
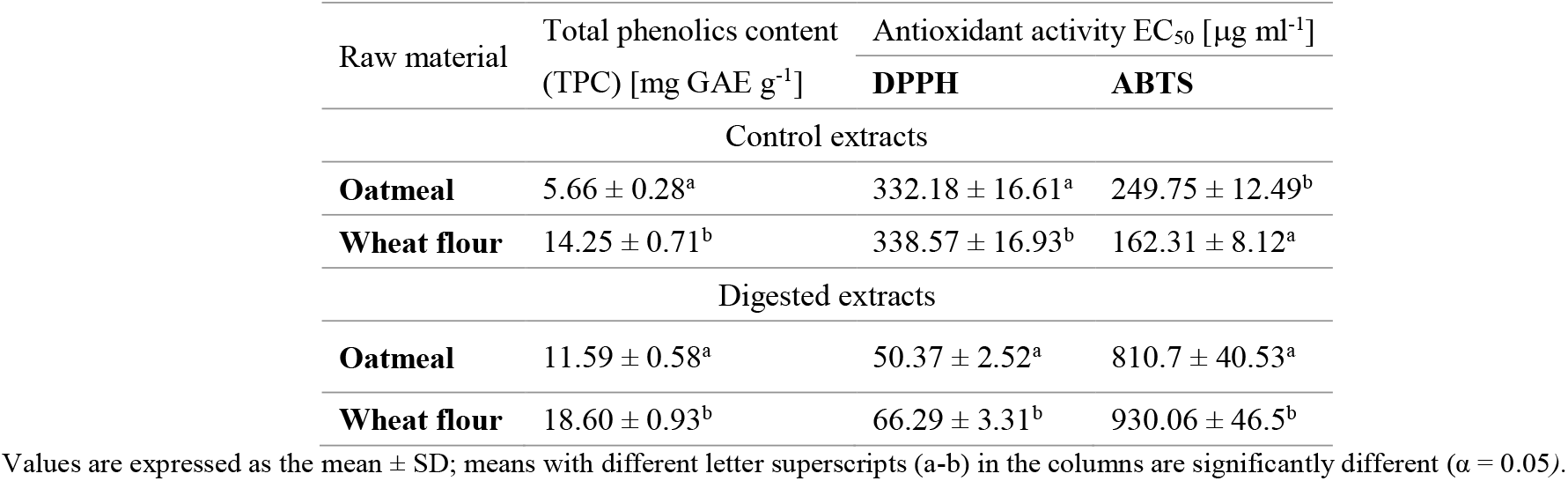
Total phenolic content and antiradical activity against DPPH ans ABTS free radicals of oatmeal and wheat flour extracts (n=9).

Generation of reactive oxygen species (ROS) during the synthesis of thyroid hormones and systemic inflammatory responses can evoke oxidative stress and damage to the thyroid gland. Consequently, its inflammation (sometimes also neoplasia) is prompted when the excessive amount of ROS is not satisfactorily managed by the cells (Ramli et al., 2017). These data highlight the need to determine the antioxidant potential of the investigated extracts. Radical scavenging assays were performed to estimate antioxidative potential of GD WF and OM extracts. They demonstrated higher antioxidative efficiency of GD OM extracts than of their GD WF counterparts, as is illustrated by EC_50_ values estimated with ABTS and DPPH assay (50.37±2.52 for OM vs. 66.29±3.31 μg DW/mL for WF). Notably, these values were considerably higher than those obtained for EtOH extracts. They also considerably differ from the data previously obtained for oat and wheat MetOH and acetonic extracts (MetOH/Oat: 510 to 18 μg/ml; (Ihsan et al., 2022), 6.57 ± 0.023 mg DW/mL acetonic/oat flour (Žilić et al., 2011). Again, the differences in ABTS results for the given product may be due to the plant species and variety, the growth conditions, the age of the plant, and the storage conditions. These discrepancies may also be attributed to the differences in the extraction methods and solvents used, which can have a significant impact on the extraction efficiency and antioxidant activity of the samples. The differences between EtOH and GD OM/WF extract activity that we obtained with ABTS and DPPH assay support this notion. As expected, the scavenging activity of the extracts was lower than that obtained for pure FA (Habza-Kowalska et al., 2021; Habza-Kowalska, Kaczor, et al., 2019). However, the data on FA content in both plants (Table 1) suggest that FA is not a primary antioxidative compound in GD OM extract. In conjunction with the differences in TPC, we show relatively high activity and potential nutritional value of OM compounds for the patients with hypothyroidism.

### 3.3. Effects of the OM and WF extracts on the activity of TPO, LOX and XO

To further assess the potential of OM compounds for the prophylactics of hypothyroidism, we investigated the influence OM and WF extracts on TPO activity (Fig. 1A-B, Table 3). Relatively strong activating effects of both GD extracts could be observed. They were corresponding to that observed for purified FA, even if its action was more efficient (Habza-Kowalska et al., 2021). Because we observed rather distinct differences in the impact of the extracts and pure FA on TPO activity, these data might indicate that *in vitro* digestion releases modulators of FA activity from the food matrix. In any case, they confirm relatively strong TPO activating effect of both extracts. Based on the data contained in Table 3, it can be assumed that the activating effect of the tested extracts consists mainly in increasing the affinity of the enzyme for the substrate (decrease in Km value), although in the case of samples obtained after *in vitro* digestion of OM, an increase in Vmax was also observed (compared to the activity of pure TPO) (Table 3). In the available literature, there are few studies on the mechanism and kinetics of enzyme activation. Shabani et al. (Shabani & Sariri, 2010) proved that same saturated and unsaturated fatty acids enhanced tyrosinase activity and affected both of kinetic parameters (decrease of Km and increase of Vmax). This type of kinetic behavior is typical of a mixed-type activator meaning that activators could bind to both the free enzyme and enzyme-substrate complex. The same situation occurred in our research in the case of GD OM samples (Fig1B, Table 3).The current literature lacks data on the mechanisms of TPO activation by bioactive compounds derived from food, so this issue requires further, comprehensive research.

**Fig. 1.**
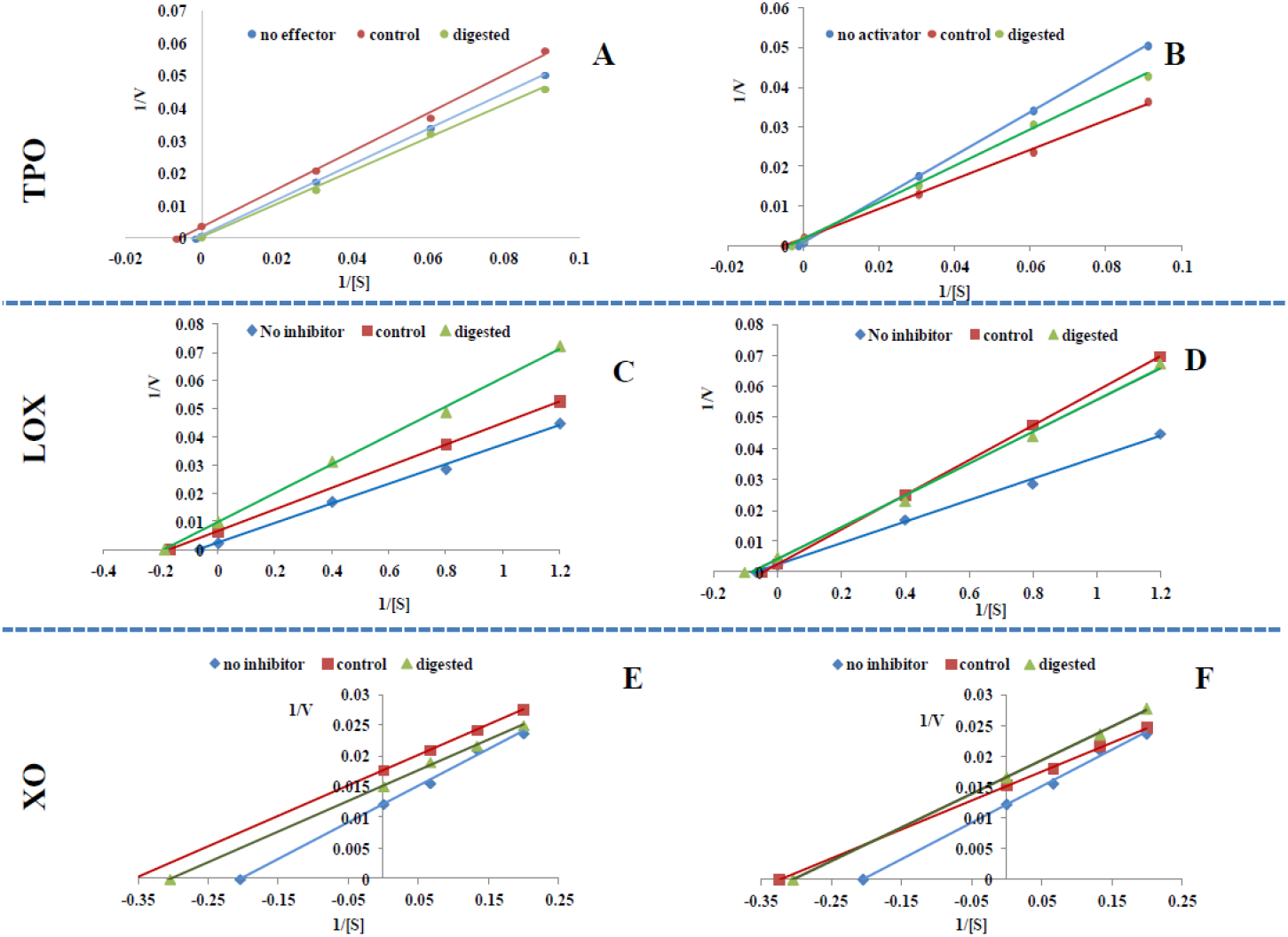
Mode of thyroid peroxidase (TPO), lipoxygenase (LOX) and xanthine oxidase (XO) affection by control and digested extracts from oatmeal (A, C, E) and wheat flour (B,D,F). Plots are expressed 1/velocity versus 1/substrate [mM] without or with extracts in a reaction solution. Guaiacol was used as a substrate for TPO, linoleic acid for LOX and xanthine for XO activity estimation.

**Table 3.**
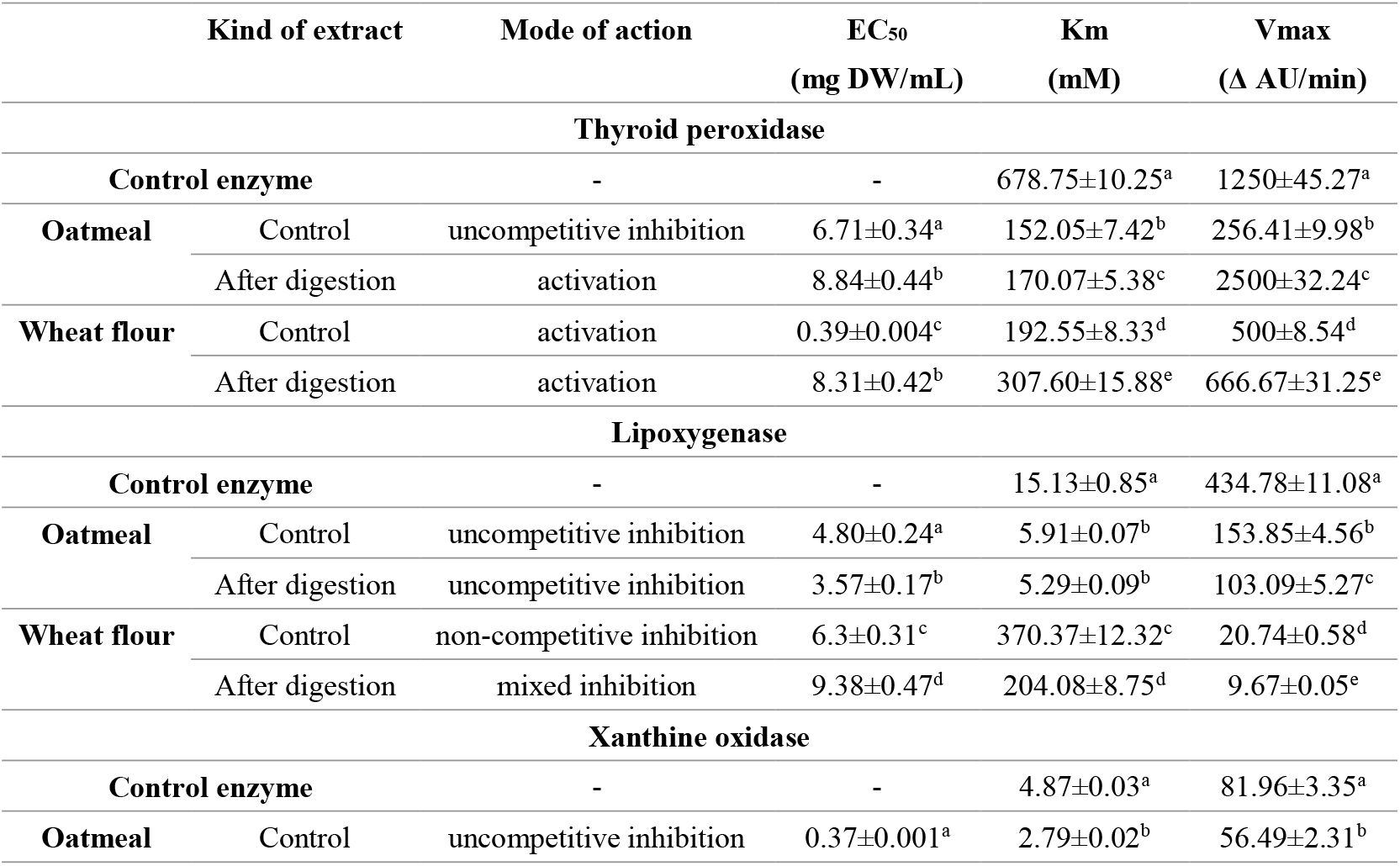
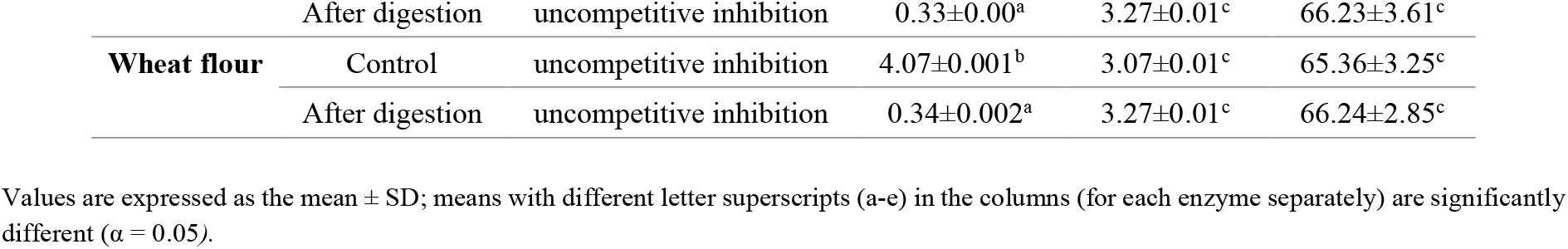
Impact of control and digested extracts from oatmeal and wheat flour on thyroid peroxidase, lipoxyhenase and xanthine oxidase activity. The EC_50_ and kinetic parameters values (n = 9)

In turn, the data shown in the Fig. 1 C-F (summarized in the Table 3) again pointed to the higher pro-health value of OM extract. Extraction-dependent differential effects of OM and WF extracts on the LOX activity are illustrated by the differences in the mode of action between WF and OM extracts and differential EC_50_. Overall, they indicate the strongest inhibitory activity of GD OM compounds on the activity of this enzyme (Table 4). Interestingly, the mode of LOX inhibition by WF extract was dependent on kind of extract; control extract acted as non-competitive inhibitor, whereas GD extract demonstrated mixed mechanism of inhibition. In the case of the OM extracts, uncompetitive mode of LOX inhibition was observed (Fig. 1C and D).

Gastrointestinal digestion also had a strong effect on the inhibitory activity of WF extracts against XO. This is illustrated by a very low EC50 values estimated for GD WF extract. In this case, however we did not observe any differences between the activities of GD OM and GD WF extract (Fig. 1 E,F; Table 3). For XO enzyme, uncompetitive inhibition was observed; The interference of wheat compounds with XO activity has already been reported. Studies of Pavia et al. (2013) demonstrated the potential of wheat bran-derived ferulic acid derivatives as XO inhibitors and scavengers of hydroxyl radical with the EC_50_ values ranging from 0.16 to 0.43 mM. Furthermore, FA derivatives had higher antioxidant activity than the parental compound (FA). Our results suggest that FA derivatives may also be responsible for the differences between OM and WF extracts. Even if this notion requires experimental verification, our study is the first to show the combined inhibitory effect of OM and WF extract on TPO, LOX and XO. It also shows the potential of OM compounds for hypothyroidism prophylactics. On the other hand, the activity of analyzed extracts (its strength and mode) non-linearly depended on their composition and concentrations. They may have serious consequences for their potential application in the prophylactics of thyroid diseases.

### 3.4. Interaction assay

When scrutinizing the bioactivity of the plant compounds, it is important to consider the interactions between food components and their effect on the bioavailability, metabolism, and on overall health outcomes of the product. Numerous reports have documented that the interactions between drug components can enhance their individual effects (Huang et al., 2019). However, our knowledge on the interactions between the modulators of pro-oxidative enzymes and TPO is limited. Our previous study and the data described above suggest the interactions between polyphenolic activators of TPO and LOX inhibitors (Habza-Kowalska et al., 2021) To assess how food matrix affects the interactions between different FA sources, we used isobolographic method, where the combination of two active substances can enhance or reduce the strength of single component influence on human health (Chou, 2006). The interaction strength is then described by the CI (Combination Index) value (Meireles et al., 2007;Chou, 2006).

The combination of an activator and an inhibitor can lead to problematic results; therefore, we did not analyze the synergistic effects of EtOH extracts on TPO activity. However, a strong synergy of stimulatory effects of GD OM and WF extracts on TPO is illustrated by relatively low CI value (0.18 ± 0.02; Fig. 2A). Similarly, strong synergy of inhibitory effects of EtOH and GD extracts on LOX activity could be seen (CI values of 0.23 ± 0.03 and 0.25 ± 0.01 respectively). Finally, we found synergistic interactions between OM and WF extracts on the inhibition of XO enzyme (CI=0.26 ± 0.04 and 0.45 ± 0.02 for EtOH and GD extracts, respectively; Fig.2 B-E).

**Figure 2.**
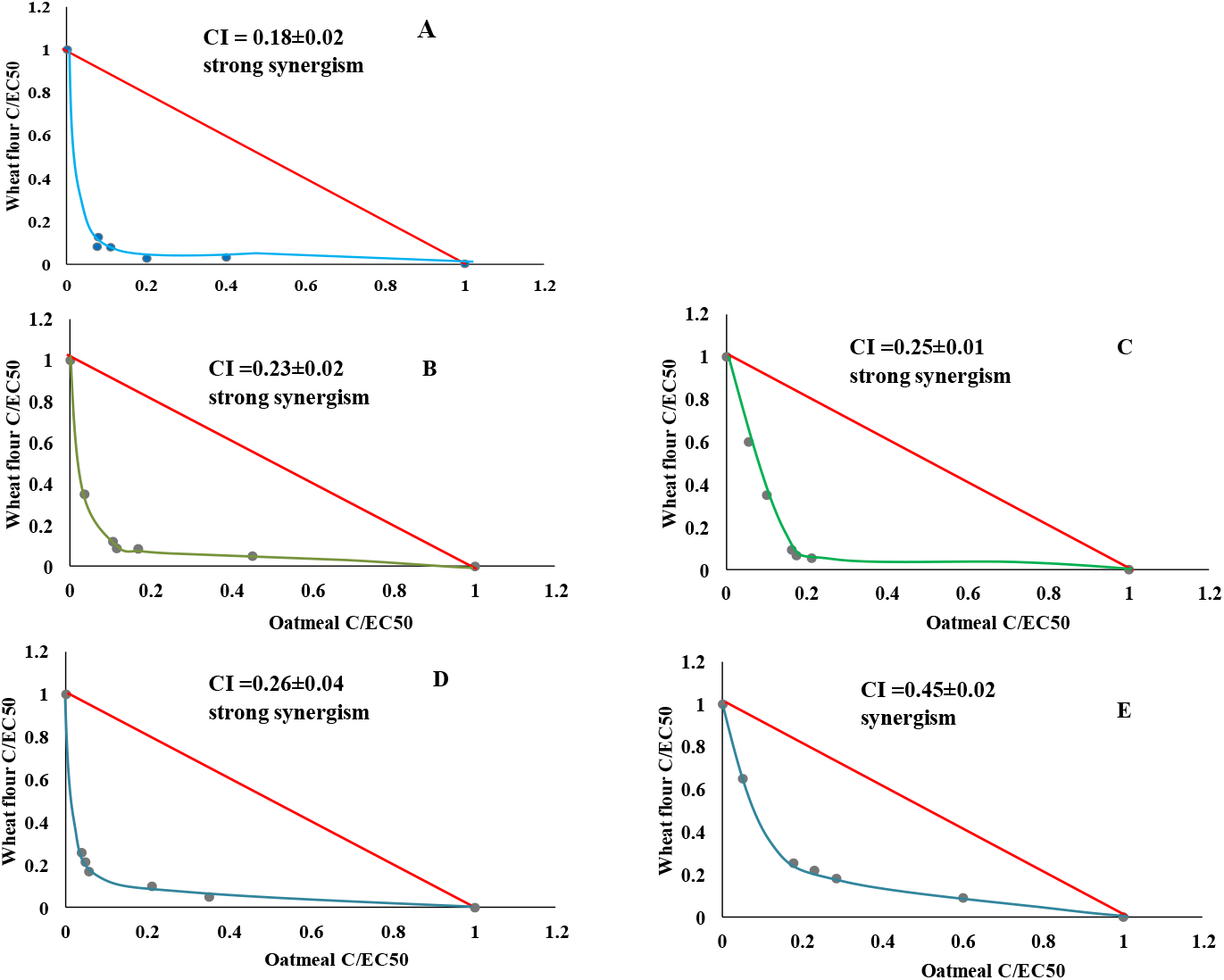
Dose – normalized isobolograms and combination index (CI) values for digested extracts of oatmeal and wheat flour with TPO activatory activity (A), control and digested extracts of oatmeal and wheat flour with LOX inhibitory activity (B and C, respectively), control and digested extracts of oatmeal and wheat flour with XO inhibitory activity (D and E, respectively).

Understanding the complex interplay between food components is crucial for a full comprehension of the impact of bioactive compounds on human health. Studies on the effect of food components on prooxidative enzymes (e.g., LOX and XO) and the analyses of the interactions between different sets of bioactive substances (incl. FA derivatives) may provide a valuable information on their potential in reducing the risk of oxidative stress and inflammation. This may be of special importance for the patients with hypothyroidism or Hashimoto disease. Our data, which show strong synergism between GD OM and WT extracts, also indicate that their phenolic content may differ, even if their basic LOX/XO inhibitory activity is similar. The same substances can affect the activity of different enzymes in diverse ways, depending on the composition of food matrix and the way of extraction. These differences may have consequences for people with thyroid diseases coexisting with oxidative stress and inflammation in the organism. On the other hand, our *in silico* observations need to be confirmed by *in vitro/in vivo* data.

### 3.5. The activity of OM anf WF extracts in vitro

#### 3.5.1. GD OM extract exerts cytostatic and pro-invasive effects in TPO^+^ B-CPAP populations

Thyroid peroxidase (TPO)^+^ B-CPAP thyroid cancer cells represent a biological model of poorly differentiated thyroid carcinomas (PDTC) that, together with anaplastic (undifferentiated) thyroid carcinomas (ATC), are associated with a poor prognosis and mainly account for thyroid cancer-related mortality. PDTC represent an intermediate stage in the progression of well-differentiated thyroid carcinoma towards ATC (K. N. Patel & Shaha, 2006). To correlate the inhibitory effects of OM and WF extracts on the activity of pro-oxidative enzymes in silico with their biological activity, we first analyzed the basic neoplastic traits of GD OM-or GD WF-treated B-CPAP cells (Fig. 3). Cell viability tests demonstrated similar dose-dependent cytotoxic activity of both GD extracts in B-CPAP model. When administered at the concentration between 0,01% to 3% (i.e. between 1.5 and 450 g of the native product, respectively), both extract reduced the fraction of viable cells to ca. 85% (Fig. 3A). In turn, a more pronounced cytostatic activity of GD OM extract (Fig. 3B) is illustrated by attenuated proliferation of B-CPAP cells in the presence of GD OM extract (to ca. 60%, compared to 75% estimated for WF extract). This promising observation confirms previous data on the bioactivity of both products (Meireles et al., 2007) and extend them to thyroid cancer. Actually, FA has long been suggested to inhibit the proliferation of cancer cells, however OM extract was less abundant in FA that WF (cf. Table 1). It suggests the involvement of other factors. Cytostatic activity of the extracts correlated with their inhibitory effects on LOX activity, which indicates that differential sensitivity of B-CPAP cells to GD OM and WF extracts can be partly related to different content of LOX modulators (incl. FA). However, it is also conceivable that the “interactome” of OM and WF compounds comprises a multitude of networked signaling pathways, which implies a necessity of further research on this topic.

**Figure 3.**
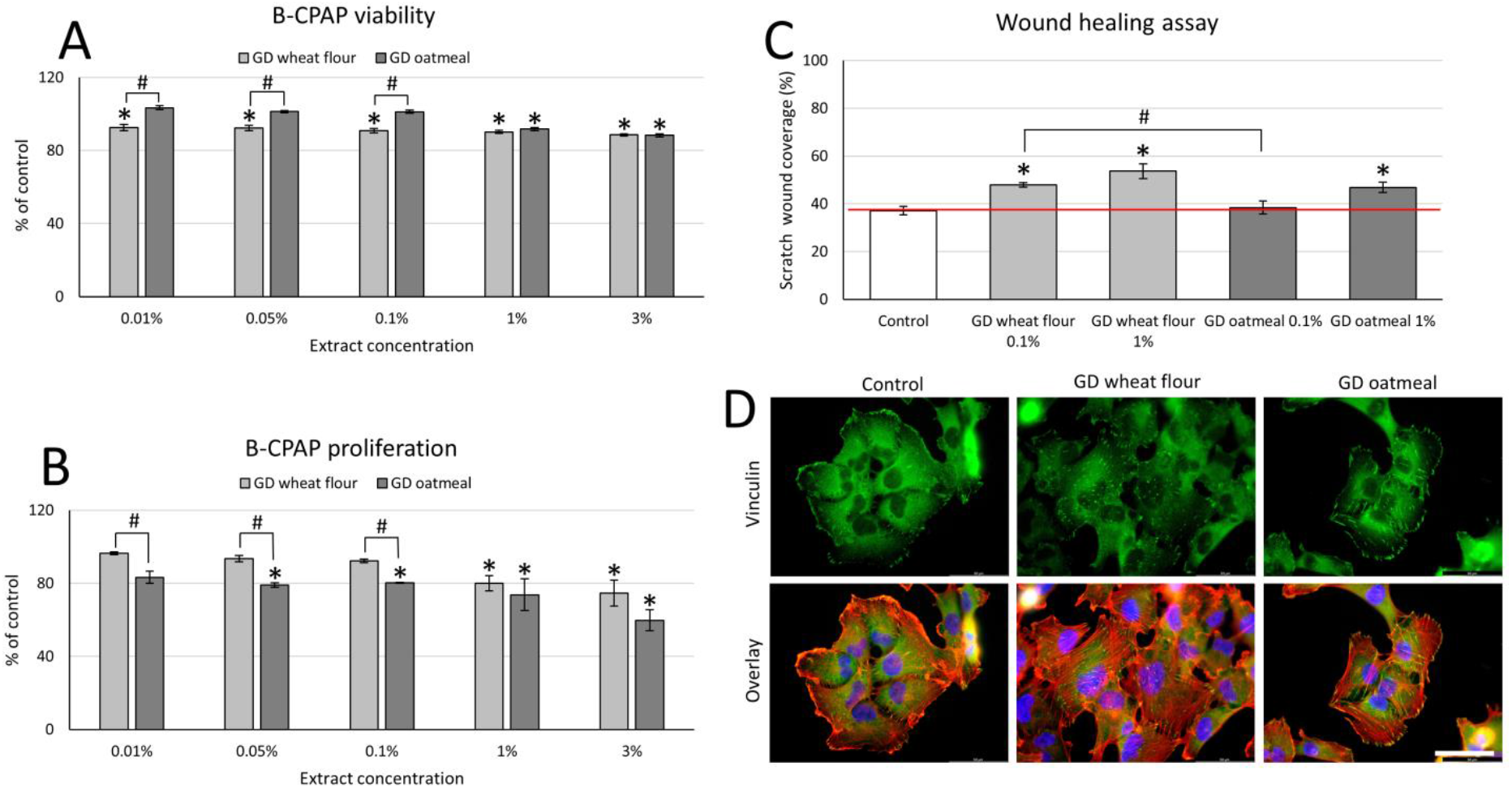
Cytostatic effects of GD WF and OM extracts in thyroid cancer B-CPAP cell populations. (A, B) B-CPAP cells were incubated in the presence of the GD extracts from WF and OM. Their viability (A) and proliferation (B) was estimated with FDA test and Coulter Counter Z2, respectively. (C, D) Effect of GD OM/WF extracts on the wound-healing efficiency (C) and actin cytoskeleton architecture/vinculin localization (D) estimated with time-lapse video microscopy and fluorescence microscopy, respectively. Statistical significance was calculated by one-way ANOVA followed by post hoc Tukey’s HSD (A,C) or Dunnet test (B), ^*,#^p<0.05 vs. relevant control. Data representative for 3 independent experiments. Bars represent SEM values. Note the cytostatic effects of the extracts, accompanied by their pro-invasive activity.

Surprisingly, the cytostatic effects of both GD extracts were accompanied by their unexpected effects on B-CPAP motility. It was slightly enhanced after their application, as illustrated by wound healing experiments (Fig. 3C). Concomitantly, we observed distinct actin cytoskeleton rearrangements (stress fibers formation) and the maturation of focal adhesions in B-CPAP cells cultivated in the presence of both extracts (Fig. 3D). Apparently, a strong cytostatic effect of WF and OM extracts applied at physiologic doses is accompanied by the induction of B-CPAP motility. We did not observe any signs of prominent apoptotic B-CPAP response, which indicates that the induction of motile phenotype rather than a selection of motile cells from heterogeneous populations accounts for the observed effects. In any case, these data may indicate the protective effect of ROS scavenging on cancer cells and/or the activation of pro-invasive signaling pathways.

#### 3.5.2. Gastrointestinal processing and anti-cancer activity of the extracts

To estimate the contribution of gastrointestinal digestion of OM and WF extracts on their activity in thyroid cancer model, we also estimated the activity of raw (PBS) extracts from both products. PBS OM and WF extracts displayed similar cytostatic activity to that of their GD counterparts (Fig. 4A,B). However PBS extract from OM had no “pro-invasive” properties (Fig. 4C). Concomitantly, we observed less pronounced actin cytoskeleton rearrangements in the cells treated with PBS extracts (Fig. 4D). Thus, gastrointestinal digestion augments the bioavailability (by release from food matrix) of protective/pro-invasive compounds of oatmeal, while retaining their cytostatic activity.

**Figure 4.**
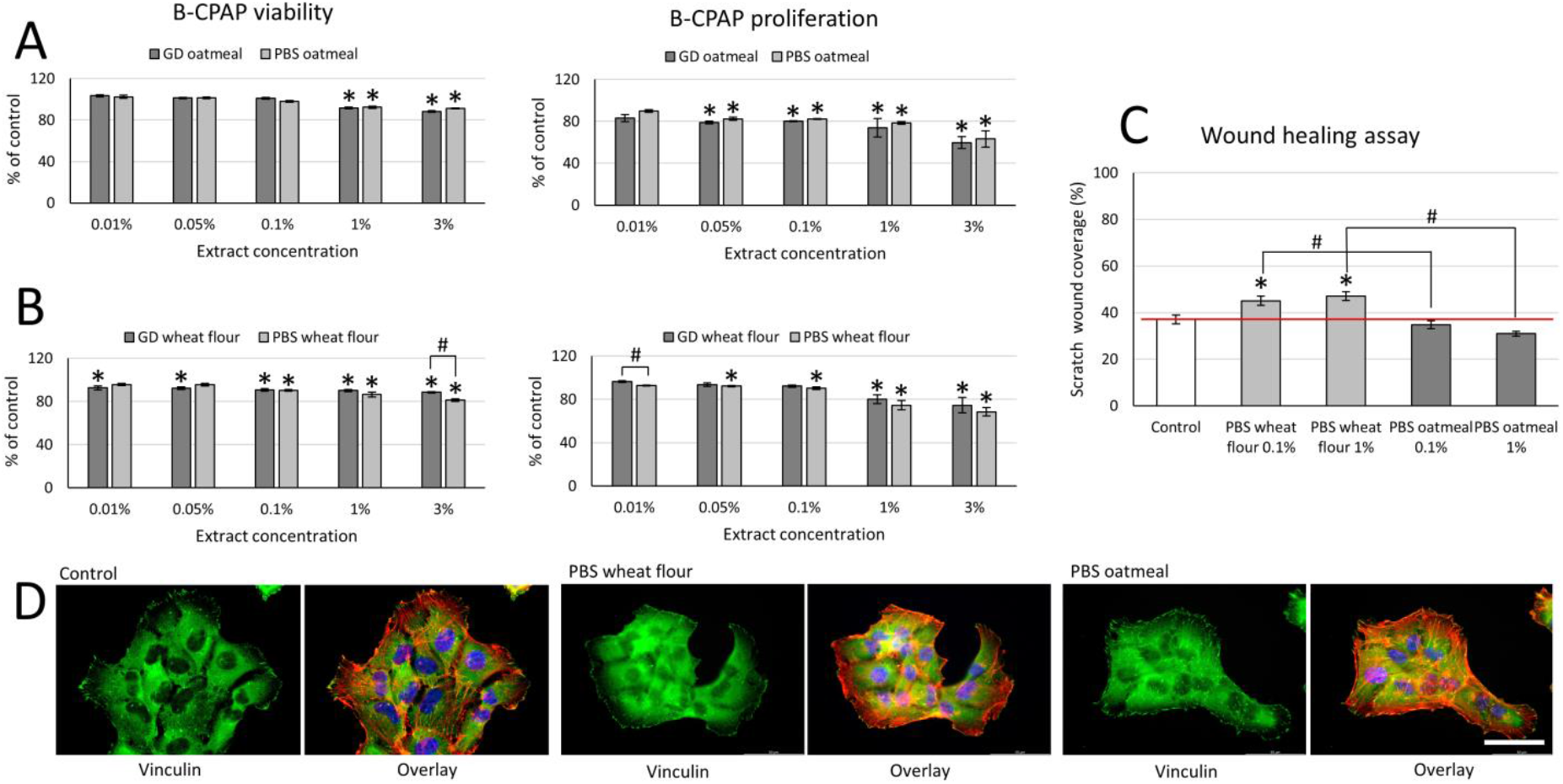
The effect of raw (PBS) extracts from wheat flour (WF) and oatmeal (OM) on the proliferation and invasiveness of B-CPAP cells. (A, B) B-CPAP viability (A) and proliferation (B) in the presence of PBS extracts from WF and OM was estimated with FDA test and Coulter Counter Z2, respectively. (C, D) Effect of PBS WF/OM extracts on the wound-healing efficiency (C) and actin cytoskeleton architecture/vinculin localization estimated with time-lapse videomicroscopy and fluorescence microscopy (GD effect = 100%). Scale bar – 50 μm. Statistical significance was calculated by one-way ANOVA followed by post hoc Tukey’s HSD (A) or Dunnet test (B, C), *,#p<0.05 vs. control. Data representative for 3 independent experiments. Bars represent SEM values. Note a less pronounced pro-invasive effects of PBS extracts.

#### 3.5.3. TPO and cell sensitivity to GD extracts

Finally, we addressed the potential role of TPO activation in the determination of thyroid cancer cell reactivity to GD OM and WF extracts. For this purpose, we used an experimental approach based on TPO^-^ (8505C) cells, which represent a well-recognized cellular model of human thyroid cancer (Meireles et al., 2007). These “dedifferentiated” cells have lost thyroid-specific markers and display higher malignancy when compared to their TPO^+^ B-CPAP counterparts. Because they do not express TPO (TPO^-^ phenotype), they are also suitable for the analyses of TPO contribution to the reactivity of thyroid cancer cells to extrinsic signals, incl. natural bio-compounds. We estimated the effect of TPO on the sensitivity of 8505C cells to both extracts by comparing their motility (wound healing) in the presence of GD/PBS OM/WF extract with the motility of B-CPAP cells in these conditions. No significant differences in the quality of 8505C and B-CPAP cells to both extracts could be seen (Fig. 5A, cf. Fig. 3C and Fig. 4C). 8505C cells remained sensitive to pro-invasive activity of 1% GD OM extract, whereas PBS extracts again displayed lower pro-invasive activity. Thus, gastrointestinal digestion increases the activity of this extract in both cellular models. Moreover, pro-invasive effects were accompanied by actin cytoskeleton rearrangements, in particular the redistribution of F-actin to cell peripheries (Fig. 5B). Collectively, a relatively high sensitivity of TPO^-^ cells to GD-released compounds indicates a marginal role of TPO activation in cellular reactivity to OM and WF extracts. Because ATC represent a subset of thyroid tumors that are associated with the worst prognosis, while TPO^-^ 8505C ATC cells display aggressive behavior and increased locoregional and distant invasion, our observations also confirm that OM/WF extracts may induce adverse effects in the patients with advanced thyroid tumors.

**Figure 5.**
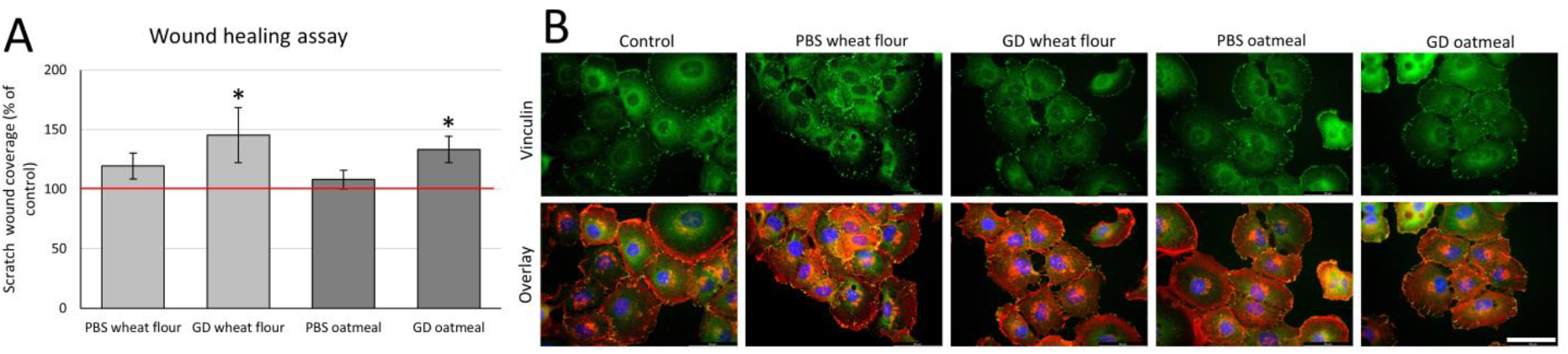
Cytostatic effects of flour and flake extracts in thyroid cancer 8505C cell populations. (A,B) Effect of GD/PBS WF/OM extracts on the wound-healing efficiency (C) and actin cytoskeleton architecture/vinculin localization estimated with time-lapse videomicroscopy and fluorescence microscopy. Statistical significance was calculated by one-way ANOVA followed by post hoc Dunnet test, *p<0.05 vs. control. Data representative for 3 independent experiments. Scale bar – 50 μm. Bars represent SEM values. Data representative for 3 independent experiments. Bars represent SEM values. Note a pro-invasive activity of both extracts in 8505C model.

Collectively, an experimental model based on TPO^+^ PDTC B-CPAP cells and their TPO^-^ ATC counterparts (8505C) cells enabled us to estimate the effect of the malignancy (differentiation status) of thyroid cancer cells’ and/or TPO expression on cellular sensitivity to plant extracts. Using this approach, we demonstrated (i) a relatively high cytostatic activity of GD OM extract, (ii) its correlation with the LOX inhibitory activity of this extract accompanied by (iii) its cytoprotective/pro-invasive effects. This points to the role of the balance between cytoprotective and cytostatic effects of compounds in the regulation of cell proliferation/motility. On the other hand, (iv) TPO activation is apparently not involved in these interactions. Cytoprotective activities of OM and WF compounds may improve cellular welfare at low concentrations. A shift from proliferation to invasion state in extract-treated cells may also be related to the “escape” strategy of stress management (Pani et al., 2010), however further research is necessary to fully elucidate mechanisms underlying this phenomenon (Pudełek et al., 2020).

## 4. Conclusions

Studies on the antiradical potential of oatmeal and wheat flour compounds (selected as a rich ferulic acid sources is based on the polyphenol database (Rothwell et al., 2013)), on their LOX/XO inhibitory activity and TPO activating effects, as well as on their effect on thyroid cancer development, provided important insights into the mechanisms underlying the health-promoting effects of these commonly consumed products. Apparently, oatmeal and wheat flour contain TPO activators and effective LOX and XO inhibitors, which can play an important role in the prophylactics of Hashimoto’s disease. To the best of our knowledge, this is the 1^st^ report on the plant products that contain such a set of bioactive compounds. Accordingly, OM and WF (especially OM!) can be used for diet supplementation in the treatment of hypothyroidism. It may have implications for the development of new functional foods and dietary supplements with multiple health benefits. However, the application of extractable hydrophilic compounds of oatmeal and wheat flour extracts as a universal supplement to interfere with thyroid cancer promotion is questionable.

Our study also provides valuable information on the effects of the gastrointestinal processing and food matrix on the bioavailability of TPO, LOX and XO modulators. In conjunction the prominent differences in the phenolic content between OM and WF extracts, we confirm that the synergy/antagonism of individual compounds is decisive for overall bioactivity of the product. Notably, the concentrations that we used in the experiments are relatively low and partly correspond to the physiologic values, which adds to the significance of our data. Therefore, our data provide new and valuable information that opens the way to further scientific investigations. It is obvious that issues related to thyroid cancer need to be confirmed in subsequent experiments. Further research is also needed to fully understand the mechanisms of the interactions between LOX/XO/TPO-related effects and cytostatic/pro-invasive activity of the tested extracts. *In vivo* and cohort studies should help to determine the safety and efficacy of these compounds as supplements or drug candidates.

## Author Contributions

Conceptualization, UGD, JC, KP. and EHK; methodology, UGD, JC, KP. and EHK .; validation, UGD, JC, KP; formal analysis, UGD, JC; investigation, EHK. KP, UGD, JC; resources, UGD, JC, KP. and EHK; data curation, EHK, UGD, KP, JC; writing—original draft preparation, EHK, KP ; writing—review and editing UGD, JC.; visualization, EHK, KP supervision, UGD, JC; project administration, UGD; funding acquisition, UGD, JC. All authors have read and agreed to the published version of the manuscript.

## Institutional Review Board Statement

Not applicable

## Informed Consent Statement

Not applicable.

## Data Availability Statement

Data available from the corresponding author on request.

## Sample Availability

Samples are available from the authors.

